# Most Beefalo cattle have no detectable bison genetic ancestry

**DOI:** 10.1101/2024.09.16.613218

**Authors:** Beth Shapiro, Jonas Oppenheimer, Michael P Heaton, Kristen L Kuhn, Richard E Green, Harvey D Blackburn, Timothy PL Smith

## Abstract

Hybridization is common among lineages in the genus *Bos*, often mediated through human management for the selection of adaptive or desirable traits. A recent example is the American Beefalo cattle breed, which was developed in the 1970s and defined as a hybrid between American bison (*Bison bison*) and cattle (*Bos taurus*). The American Beefalo Association (ABA) typically require ⅜ bison ancestry to qualify as Beefalo. Here, we sought to characterize admixed ancestry among Beefalo as a component of a larger project to understand the role of hybridization in shaping present-day diversity in bison and cattle. We generated genomic data from 50 historical and present-day Beefalo and bison hybrids, including several important founding animals, as well as from 10 bison originating from commercial herds that represent potential sources of bison ancestry in Beefalo. We found that most Beefalo did not contain detectable bison ancestry. No individual Beefalo within our data set satisfies the ancestry requirements specified by the ABA, although several Beefalo had smaller proportions of bison ancestry (2-18%). Some beefalo had detectable zebu cattle ancestry (2-38%), suggesting that hybridization of taurine and zebu cattle may contribute to morphological similarity between some Beefalo and bison. Overall, ancestry profiles of Beefalo and bison hybrid genomes are consistent with repeated backcrossing to either parental species rather than the breeding between hybrids themselves, implying significant barriers to gene flow between bison and cattle. Our results call into question the ⅜ bison ancestry targeted by the breed association and demonstrate the value of genomic information in examining claims of interspecies gene flow among *Bos* species.

## Introduction

Gene flow has been common among lineages within the genus *Bos*, including among cattle (*Bos taurus*), bison (*Bison bison*), yak (*Bos mutus*), and gaur (*Bos gaurus*) (Wu et al. 2018; Zhang et al. 2020). Much of this gene flow was facilitated by human livestock management and breeding. For example, hybridization led to gene flow between taurine (*Bos taurus*) and zebu (*Bos indicus*) cattle breeds in the Near East during the Bronze Age (Verdugo et al. 2019), and later in Africa (Kim et al. 2020) and North America (McTavish et al. 2013). Human-mediated gene flow also occurred between more deeply diverged *Bos* lineages, including Chinese cattle and banteng (Chen et al. 2018) and yak and Tibetan cattle (Wu et al. 2018). In each case, gene flow is believed to have conferred genetic benefits that led to easier management of hybrid herds or enabled adaptation to local environmental conditions (Zhang et al. 2020).

While admixture is common among *Bos* lineages, genomic incompatibilities often induce reproductive challenges to the hybrids. These incompatibilities are probably due to the considerable divergences across the *Bos* clade, with the oldest split between cattle and the other lineages dating to around three million years ago (Wang et al. 2018). Except for hybrids of taurine and zebu cattle, hybrids of all other *Bos* species follow Haldane’s rule, in which male offspring of F1 crosses are sterile (Zhang et al. 2020). While taurine-zebu cattle hybrids are fertile, hybrid African cattle have uneven proportions of zebu nuclear and mitochondrial ancestry (Kwon et al. 2022; Ward et al. 2022), which may be due to mitonuclear incompatibilities. Analysis of the dynamics of speciation and recurrent gene flow among *Bos* would improve understanding of the evolutionary process of species divergence.

Beefalo are a purported example of interspecies *Bos* hybridization. The American Beefalo Association (ABA) defines Beefalo as a stable hybrid cross with ⅝ cattle and ⅜ bison ancestry (“American Beefalo Association”). The Beefalo breed was established in the early 1970s by Mr. D.C. “Bud” Basolo through complex interbreeding of the two species (Miller 1982). The establishment of this breed followed a long history of attempts to crossbreed bison and cattle for commercial production (Boyd 1908; Goodnight 1914). As early 20th century breeders had been unable to create a viable hybrid population due to reproductive difficulties (Deakin et al. 1942), the establishment of the Beefalo breed was both surprising and celebrated (Miller 1982; “Business” 1973).

Basolo did not reveal the pedigree for his original Beefalo, but subsequent analyses have allowed better understanding of the breed’s origins. Paim et al. (2020) illustrated two pedigree approaches that could create a ⅜ – ⅝^th^ combination, both of which require first generation hybrid males to be fertile. Analyses of Beefalo karyotypes showed that all Beefalo possess taurine Y chromosomes (Lenoir and Lichtenberger 1978), such that the initial cross would have to have been a bison cow and a taurine bull. Intriguingly, Stormont et al. used blood typing to show that foundational Beefalo almost completely lack bison-specific markers (Stormont et al. 1986). Beefalo ancestry has not, however, been evaluated with genomic information, though the breed was granted a special roll stamp for voluntary federal inspection by the United States Department of Agriculture (USDA, 1984, 9 CFR part 352) and the ongoing sale and purchase of pedigreed animals.

Here, we present results of the first genome-wide investigation of the ancestry composition of the Beefalo breed. We generate whole-genome shotgun (WGS) data from 47 Beefalo, three bison hybrids reported to carry majority bison ancestry, and ten commercial bison that represent a potential source of bison ancestry in Beefalo, and co-analyze this data along with publicly available genomes from a range of bovids.

These samples were donated to the USDA Agricultural Research Service (ARS) National Animal Germplasm Program (NAGP) collection and include ancestors from the early establishment of the breed, such as Joe’s Pride, a foundational Beefalo bull that Mr. Basolo sold to a Canadian breeder’s firm for $2.5 million in 1975 (“Most Expensive Cattle”). We find that most Beefalo, including key founding individuals, contain no detectable bison ancestry, while bison hybrids have approximately their reported bison ancestry. Zebu ancestry is common among Beefalo, suggesting that breeding with zebu cattle, which share a number of phenotypic similarities with bison in commercially-relevant traits (Zeder 2006), was an early strategy in the establishment of the Beefalo breed. Finally, we find that some Beefalo contain small amounts of bison ancestry, ranging from 2-18%. While this is less bison ancestry than the targeted 37.5%, these genomes prove that bison and cattle can hybridize in some circumstances. However, we always infer minor parental ancestry to be in a heterozygous state, providing evidence only of backcrossing to either parent, rather than of breeding between hybrids themselves. This is inconsistent with a scenario in which a stable bison-cattle hybrid Beefalo population had been established.

## Results

### Sequencing Beefalo genomes

We generated genomic data from 47 registered Beefalo and three bison hybrids using preserved semen samples. Beefalo semen samples were obtained from the USDA, ARS, NAGP (Supplementary Table 1). Most of the NAGP Beefalo samples were collected during the 1970s and 80s, around the time when the breed was founded, and donated to NAGP around 2007. We retrieved bison ancestry composition for each sample from metadata accompanying NAGP samples. These 50 Beefalo and bison hybrids had reported bison ancestry ranging from 75% in a bison hybrid to 12.5% in several Beefalo, with the bison ancestry of most Beefalo reported as the breed-standard 37.5%. We sequenced each sample to 2.7x median genomic coverage (range: 1.1-39x; Supplementary Table 1). Seven prominent Beefalo were selected for higher coverage sequencing (>30x), including animals from the Basolo foundation herds, from which Beefalo initially originated, and others with different proportions of reported bison ancestry.

We generated high coverage (>17.5x) genomes of ten bison from commercial herds (Supplementary Table 1). We added these to a larger data set of previously published bison genomes (Yang et al. 2020; Stroupe et al. 2022; Wu et al. 2018; Shirazi et al.2022) and genomes from several breeds of zebu and taurine cattle (Heaton et al. 2016); Supplementary Table 1), as well as the genome of Buzz, an F1 Yellowstone bison-Simmental cattle hybrid (Oppenheimer et al. 2021). Our final data set included 41 bison, 26 cattle, and 51 Beefalo and bison hybrid genomes. For some analyses we also incorporated published genomes from outgroups, including yak (Qiu et al. 2012, 2015), gaur (Heaton et al. 2016), banteng (Heaton et al. 2016), and water buffalo (Sun et al. 2020). Pseudohaploid genotypes were then called on 5.29M biallelic autosomal variants ascertained in gaur, an outgroup to bison and cattle, for downstream analyses.

### Estimating bison ancestry in Beefalo

Most Beefalo samples cluster closely with taurine cattle in principal component analysis (PCA) conducted on bison and both cattle subspecies, indicating they do not contain appreciable bison ancestry (Fig. 1A). This includes several foundational individuals, such as Joe’s Pride (NAGP5887). The first principal component (PC1) separates bison from cattle, with the F1 hybrid Buzz falling halfway along this axis, while PC2 distinguishes taurine and zebu cattle. While Beefalo samples group with taurine cattle, the three bison hybrids (>50% bison ancestry) fall in an intermediate position between bison and cattle (Fig. 1A). These individuals fall closer to bison than does Buzz, confirming their majority bison ancestry. Eight Beefalo are slightly shifted toward bison in the PCA, demonstrating genetic affinity with bison, though the position of these individuals on PC1 suggests they are unlikely to contain the breed-standard 37.5% bison ancestry (Figs. 1B, C). Beefalo formed a cline between taurine and zebu cattle on PC2, suggesting zebu ancestry is a widespread and variable ancestry component found in Beefalo.

**Figure 1.**
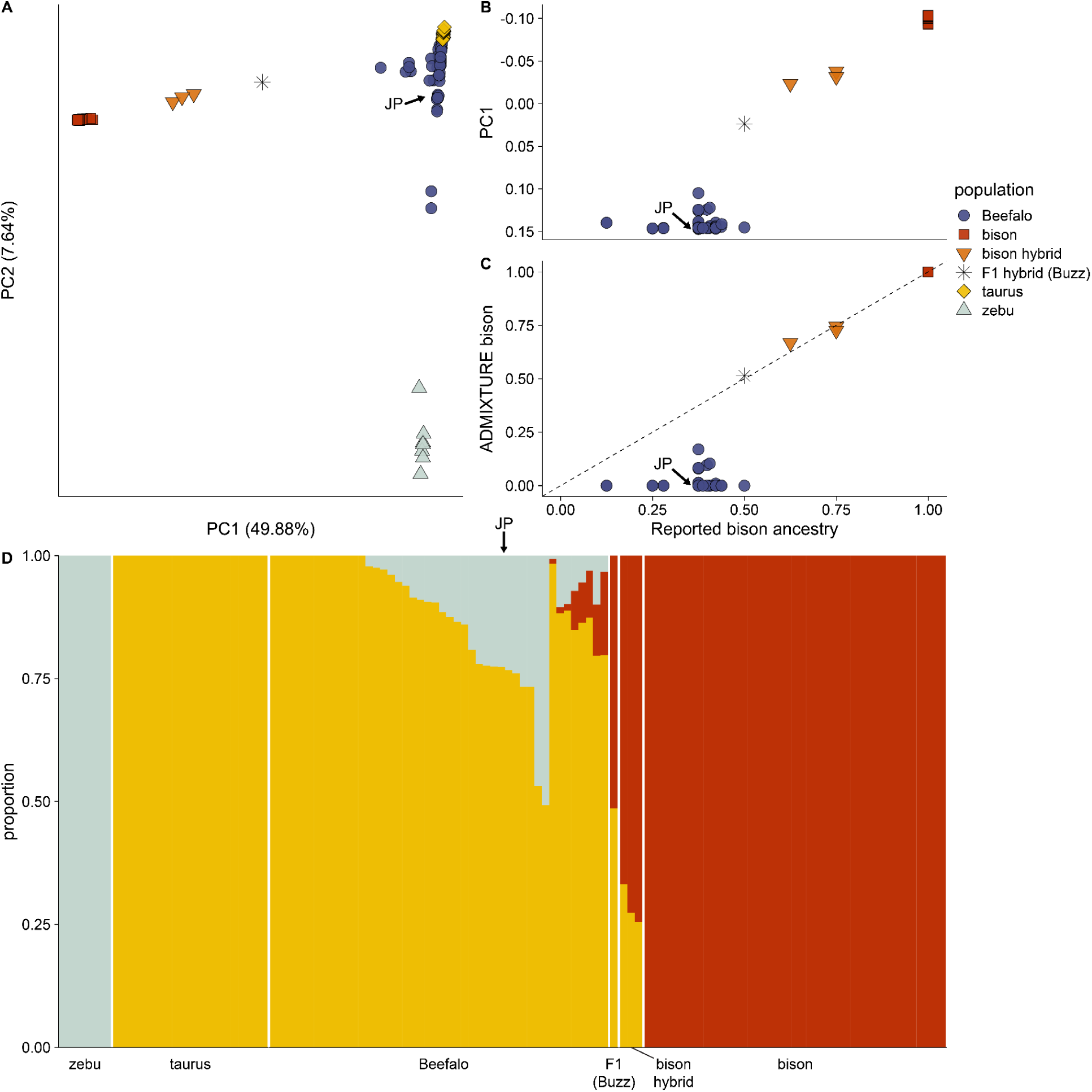
**A)** Principal component analysis with Beefalo (blue), bison-cattle hybrids (orange), and a bison-cattle F1 hybrid (black star) projected onto axes computed from bison (red) and taurine (yellow) and zebu (gray) cattle. PC1 separates bison and cattle while PC2 separates the two cattle groups. **B)** Comparison of Beefalo reported bison ancestry and position on PC1. Placement of bison hybrids on PC1 is correlated with reported bison ancestry, though Beefalo do not follow this trend. **C)** depicts the same as in **B)** though compares inferred bison proportions from ADMIXTURE to reported bison ancestry. Reported and inferred bison proportions match well for bison hybrids, but all Beefalo were inferred to have less bison ancestry than reported. **D)** Autosomal ADMIXTURE analysis of bison, taurine and zebu cattle, bison hybrids, and Beefalo. The position of Joe’s Pride (JP) is depicted with an arrow in **A-D**.

One animal labeled as Beefalo in the NAGP database, NAGP9109, fell within the zebu cluster and is likely of 100% zebu origin. The reported pedigree in the NAGP for this animal lists its composition as ½ Brahman, ¼ Charolais, ⅛ bison, 1/16th Hereford, and 1/16th Shorthorn, but the American Brahman Breeders Association records this animal (#309519) as purebred Brahman, which is a zebu breed (5 of the other 6 zebu individuals analyzed here are Brahman cattle). We infer that NAGP9109 was erroneously labeled as Beefalo by the contributors. This result highlights that PCA analyses of even low coverage genomes can uncover inconsistencies between genomic ancestry and animal metadata.

An unsupervised ADMIXTURE analysis supports the conclusion that the majority of Beefalo have no bison ancestry. This analysis estimates the ancestry profiles present across all bison, cattle, bison hybrid, and Beefalo individuals examined for a given value of *K* source populations. At *K* = 3, bison, taurine cattle, and zebu cattle are divided into homogenous groups, with Beefalo assigned variable levels of these three ancestry types, mirroring the major patterns seen in the PCA (Fig. 1D). As in the PCA, the three bison-cattle hybrid backcrosses are inferred to have majority bison ancestry, consistent with their pedigrees, while 39 out of the 47 sampled Beefalo, including Basolo founding individuals, possess no bison ancestry. The remaining eight Beefalo tested are estimated to have a small amount (<18%) of bison ancestry, although less than was indicated by pedigrees and below the breed standard defined by the ABA (Fig. 1C).

These eight individuals are those shifted toward bison along PC1 (Figs. 1A, B). The unsupervised ADMIXTURE analysis correctly assigned the F1 bison-Simmental genome equal amounts of taurine and bison ancestry, demonstrating that this analysis is sensitive to the presence of large fractions of bison ancestry. Similar estimates of bison ancestry are observed at other values of *K* (Supplementary Fig. S1), which supports the modeling of Beefalo ancestry as coming from three main components, corresponding to taurine and zebu cattle, and bison (Supplementary Fig. S2). Supervised ADMIXTURE clustering, which models Beefalo ancestry as coming specifically from bison, taurine, and zebu source panels, provides similar estimates to those obtained from the unsupervised ADMIXTURE analysis (Supplementary Fig. S3).

Examining allele sharing between Beefalo and bison relative to taurine cattle confirmed the results from PCA and ADMIXTURE, underscoring that 39 out of 47 examined Beefalo have no appreciable bison ancestry, while those that do have far less than their reported amounts. *D*-statistics of the form *D(Beefalo, taurus; bison, water buffalo)* again show that 39 Beefalo have no excess affinity with bison compared to taurine cattle (−13.04 < *Z <* 3.14), although the same eight Beefalo identified in PCA and ADMIXTURE as having bison ancestry also have an excess of bison alleles (6.16 < *Z <* 34.86), confirming their bison ancestry (Fig. 2A). f4-ratios estimate that these Beefalo derive between 2-18% of their genomes from bison (Fig. 2B), consistent with estimates from ADMIXTURE and less than their reported bison ancestry, which ranged from 37.5-50%. *D*-statistics also confirm that 18 Beefalo individuals have significantly more zebu alleles compared to taurine cattle (3.64 < *Z <* 21.03), demonstrating that zebu ancestry is widespread across Beefalo (Fig. 2C). These 18 individuals with excess zebu affinity had no evidence of bison ancestry using *D*-statistics, suggesting that allele sharing with bison, which are deeply diverged from both taurine and zebu cattle, does not complicate the inference of zebu ancestry in these individuals. Allele sharing results are similar using yak as an outgroup, instead of water buffalo (Supplementary Fig. S4).

**Figure 2.**
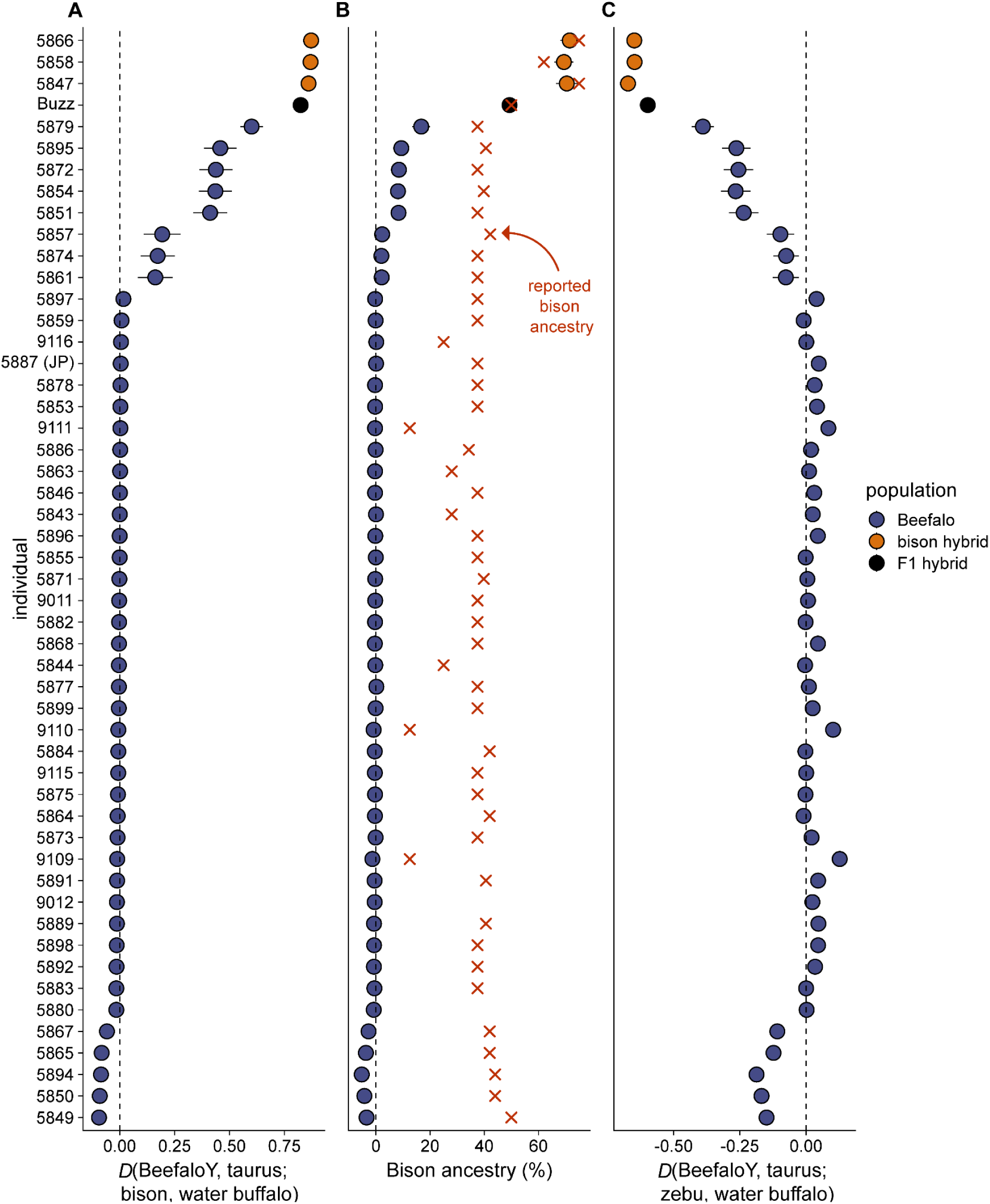
Allele sharing statistics using individual bison hybrids and Beefalo. Individuals are shown on the Y axes. Beefalo are shown in blue and bison-cattle hybrids in orange, with a bison-cattle F1 shown in black. JP = Joe’s Pride. **A)** Testing for the presence of bison ancestry using statistics of the form D(Beefalo, taurine cattle; bison, water buffalo). **B)** Quantification of the proportion of bison ancestry using f4-ratios. All bison hybrids have at least 50% bison ancestry, while only 8 Beefalo are inferred to have bison ancestry, ranging from 2-18%. Reported bison ancestry is shown in red. **C)** Testing for allele sharing between Beefalo and zebu cattle, relative to other taurine cattle. Many individual Beefalo are inferred to have significant allele sharing with zebu cattle. All Beefalo which were inferred to have bison ancestry in panels **A)** and **B)** display significantly negative values (due to the presence of bison ancestry, which is an outgroup to both cattle breeds), showing that any inferred zebu ancestry is not due to the presence of bison ancestry. For all panels, error bars depict 3 standard deviations.

Local ancestry inference across individual Beefalo and bison-cattle hybrid genomes provides similar estimates of overall Beefalo ancestry, inferring an absence of bison ancestry across the 37 Beefalo that lacked evidence for such ancestry in previous analyses (Fig. 3). Three bison hybrids are inferred to have ∼75% bison ancestry, while eight Beefalo have detectable bison ancestry, ranging from 2-18%. In the 8 Beefalo with bison ancestry, that ancestry tends to be present in large contiguous blocks, often tens of megabases in size, indicative of recent admixture (Fig. 3A,B). Bison ancestry in Beefalo is always found in a heterozygous state, consistent with a scenario in which these individuals are the product of repeated backcrossing to cattle of an initial F1 hybrid. Thirty-eight Beefalo also have zebu ancestry (Fig. 3D) at variable levels ranging from 2-38%, with all but two Beefalo having between 2-18%. As with bison ancestry, zebu ancestry is found in large blocks, but with many smaller blocks than seen with bison ancestry, suggesting an earlier date for admixture (Fig. 3C).

**Figure 3.**
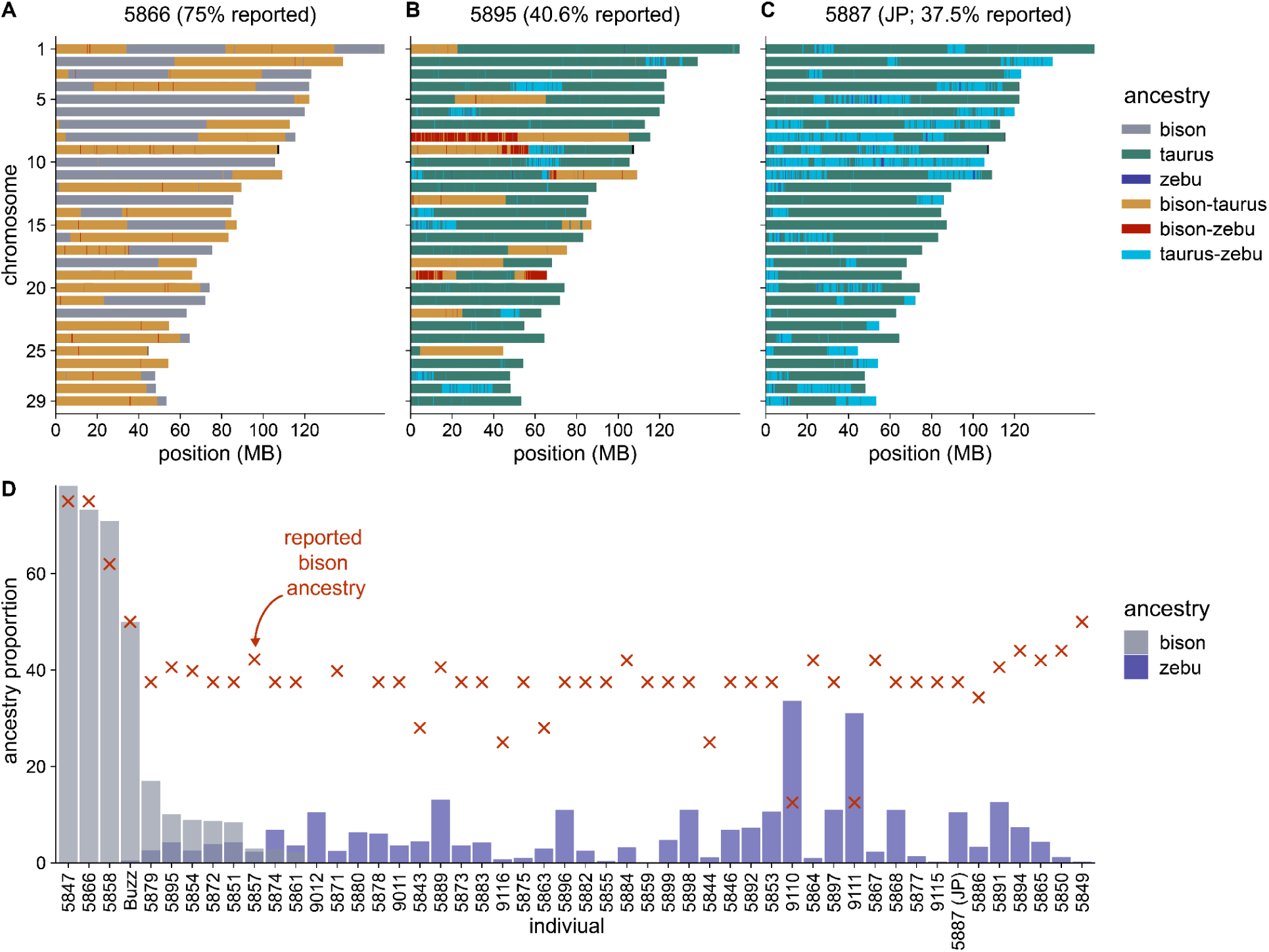
Local ancestry inference of Beefalo and bison hybrid genomes, using bison and taurine and zebu cattle as potential sources. Inferred ancestry across the all autosomes are shown for **A**) a bison hybrid with reported 75% bison ancestry, **B**) a Beefalo with 37% reported, and 12% detected, bison ancestry, and **C**) Joe’s Pride, a foundational Beefalo with 40.6% reported, and no detected, bison ancestry. Ancestry along the genome is colored by inferred source: homozygous bison, taurine, and zebu ancestry are shown in gray, green, and blue, respectively, with heterozygous bison-taurine, bison-zebu, and taurine-zebu ancestry shown in yellow, red, and light blue. Panel **D**) shows the inferred global bison and zebu ancestry proportions for each bison hybrid and Beefalo individual, with bison ancestry proportion shown in gray and zebu proportion shown in blue.

Several lines of evidence attest to the efficacy of using this local ancestry inference approach for modeling Beefalo ancestry. The F1 bison-taurine cattle hybrid was inferred to be almost entirely heterozygous for bison and taurine cattle across the genome (Supplementary Fig. S5), demonstrating the sensitivity of this approach to distinguishing diploid ancestry states between these groups. Further, the method assigned largely uniform and correct ancestry proportions to individual bison and cattle that were not part of the source panels, while Brangus cattle, a known taurine-zebu hybrid breed, were assigned large fractions of ancestry to both cattle groups in expected amounts (Supplementary Fig. S6). For the seven Beefalo sequenced to high coverage, we also modeled ancestry using downsampled read counts and called genotypes, which yielded nearly identical results (Supplementary Fig. S7).

### Beefalo sex chromosomal ancestry

Analyses of sex chromosomes reveal that the main mechanism for introducing bison ancestry into Beefalo was breeding with bison bulls. Beefalo with evidence of bison ancestry have less bison ancestry on X chromosomes compared to autosomes (Fig. 4A,C). Further, all Beefalo have Y chromosomes associated with taurine cattle, while the bison hybrid backcrosses have bison Y chromosomes (Fig. 4B). These patterns are the opposite of what would be expected if bison ancestry were introduced maternally, which would result in an excess of bison rather than cattle ancestry on the X chromosome (Lenoir and Lichtenberger 1978). Unfortunately, as semen was the source of genomic DNA, we could not assemble mitochondrial genomes to assess whether the Beefalo maternal line would provide any additional insight into these patterns.

**Figure 4.**
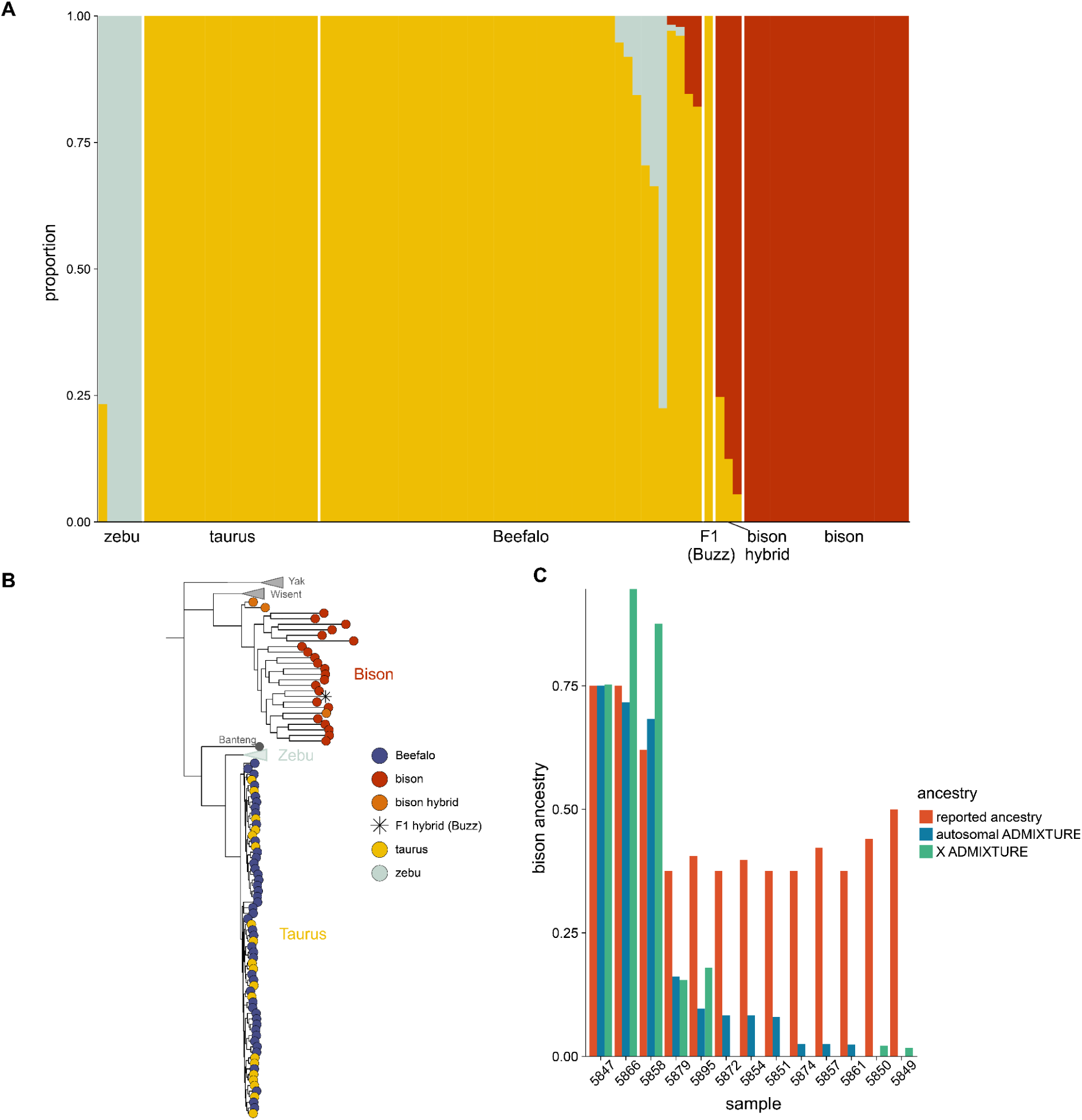
**A)** ADMIXTURE analysis of bison, taurine and zebu cattle, bison hybrid, and Beefalo X chromosomes. Beefalo are largely inferred to have X chromosomes derived entirely from taurine cattle, though variable amounts of zebu and bison ancestry are also present in some Beefalo. The position of Joe’s Pride (JP) is indicated with an arrow. Bison hybrids have majority bison ancestry. **B)** Bovid Y chromosomal phylogeny. The zebu clade is collapsed, as are those containing yak and wisent. Water buffalo (not shown) was used as an outgroup. Beefalo all have taurine Y chromosomes, while bison hybrids have bison Y chromosomes. **C)** Comparison of the reported (red) and ADMIXTURE inferred bison ancestry proportion on the autosomes (blue) and X chromosome (green) for the three bison hybrids and eight bison which had detectable autosomal bison ancestry.

## Discussion

Only eight of the 47 Beefalo that we sampled contained detectable bison ancestry, and those eight had substantially less (2-18%) than the 37.5% specified as the breed standard set by the ABA. Our sampling represents a complete survey of individuals in the USDA NAGP biobank, and includes important foundational animals that were involved in establishing Beefalo, such as those from original Basolo herds. Notably, important foundational individuals, including Joe’s Pride (NAGP5887), which was sold for $2.5 million to a Canadian breeders group (“Most Expensive Cattle”), lack bison ancestry. While these results show that interbreeding between bison and cattle is possible, they also prove bison ancestry encompasses a much smaller portion of the Beefalo breed than has been claimed.

Our finding of little to no bison ancestry in founding Beefalo individuals is aligned with challenges hybridizing bison and cattle (Goodnight 1914; Murdoch 2018). Artificial insemination of taurine cows with bison semen results in 77% calf mortality and sterile male calves (Anstey 1986). In fact, Canadian government research reported that no functional males carrying more than 12.5% bison ancestry were ever observed (Brower 2008). In Beefalo individuals with bison ancestry, such ancestry was found exclusively in a heterozygous state, implying that repeated backcrossing to parental species, rather than breeding of hybrids themselves. A depletion of bison ancestry on the X chromosome and the presence of taurine Y chromosomes in all Beefalo, with bison hybrids exclusively having bison Y haplotypes, is consistent with this scenario. Therefore, it seems that extensive reproductive barriers exist to establishing a hybrid bison-cattle population.

While bison ancestry was surprisingly underrepresented in our Beefalo sample, the majority of the Beefalo genomes we sequenced contained some zebu cattle ancestry, suggesting zebu/taurine interbreeding may have been used as a mechanism to manipulate the Beefalo phenotype. Zebu cattle are known for heat and drought tolerance (Kumar et al. 2016; Vajana et al. 2018; Hansen 2004), lower nutritional demands relative to taurine cattle (Hunter and Siebert 1985; Hennessy, Williamson, and Darnell 2000) and their humped appearance (Heath 1979), and so share a number of desirable attributes with bison. The Beefalo breed standard does not require cattle ancestry to be of taurine origin, though if early breeders intentionally incorporated zebu ancestry while creating the breed, they left this detail out of the animals’ reported pedigrees. For example, the Basolo founder individual Joe’s Pride has a recorded pedigree that attests Hereford, Charolais, and bison (37.5%) ancestry, but we estimated that he had ∼10.5% zebu ancestry.

This study is the first examination of Beefalo ancestry using whole genome data, but it is not the first to question claims surrounding bison ancestry in the breed. In an investigation of the paternal origins of Beefalo, Lenoir and Lichtenberger (1978) used karyotyping to determine that all 12 Beefalo bulls they examined had cattle Y chromosomes, in concordance with our results. While there are plausible scenarios for Beefalo with 37.5% bison ancestry to have cattle Y chromosomes, these require early-generation crosses to be fertile. In a later study, Stormont et al. (1986) used species-specific blood group markers to show that only one of the 148 Basolo Beefalo they tested had any alleles found predominantly in bison. However, they also note that some later Beefalo, separate from the initial founding herds, did display some bison-associated markers, perhaps agreeing with our finding of a small fraction of Beefalo possessing appreciable bison ancestry. Blood typing provides limited insight into ancestry proportions, however, demonstrating the utility of genomic information in validating specific breeding claims.

Our genome-wide analyses were primarily conducted on ∼2x coverage genomes with Beefalo samples obtained from the USDA NAGP, but we believe our Beefalo ancestry estimates are robust and representative of the breed at large. We derive highly concordant ancestry estimates across a range of approaches, including ADMIXTURE, f4-ratios, and local ancestry inference techniques, and for animals whose ancestry is known, such as an F1 bison-cattle hybrid, these methods assign the correct ancestry proportions. Additionally, we derive similar ancestry estimates in high coverage (30-42x) and downsampled data for the seven individuals we sequenced more deeply. We did not sample present-day sources of Beefalo exhaustively, instead choosing to focus largely on animals that originate at or near the founding of the breed. Beefalo herds are generally established by breeding with fullblood (37.5% bison) Beefalo individuals, rather than through backcrosses with bison (“American Beefalo Association”), so surveying breed founders is an effective way of documenting the bison ancestry across the breed. However, a larger sample of Beefalo across current producers would provide a more comprehensive understanding of the breadth and variation of bison ancestry in Beefalo.

The extent of gene flow and the existence of reproductive barriers among *Bos* species remain incompletely explored. Breeders worked throughout the 20th century to create commercially-viable bison-cattle hybrids using bulls and cows with different ancestries. All of these efforts, including Beefalo, failed to create a stable hybrid population, implying that considerable barriers to gene flow exist between these two species.

Though genomic incompatibilities among bison and cattle are hinted at by these breeding efforts themselves, which reported sterility in male hybrids even after backcrossing to cattle for several generations (Deakin et al. 1942; Brower 2008), they have not been detected directly. The general absence of bison ancestry in Beefalo also calls attention to several other examples of assumed gene flow among *Bos* species that have yet to be fully characterized. This includes introgression from cattle into wisent (Wecek et al. 2017; Soubrier et al. 2016) and American bison, the latter of which has been suggested to be widespread over the past two centuries and has led to the presence of cattle ancestry in all bison today (Stroupe et al. 2022; Halbert and Derr 2007), though has not been fully examined with genomic evidence.

## Materials and Methods

### Genomic data collection

Beefalo semen samples (n = 47) were obtained from the USDA, ARS, NAGP collection at Ft. Collins Colorado in the USA (Supplementary Table 1). The three bison hybrid semen samples were also obtained from the NAGP. The majority of the NAGP samples were collected in the 1970s and 1980s, stored by breeders, and donated to NAGP circa 2007 as a geographically diverse set of Beefalo. Purebred bison meat samples (n = 10, tongue) were purchased from commercial processors and represent animals from three different commercial bison herds (Supplementary Table 1).

DNA was extracted from semen samples with a modified standard phenol-chloroform method. Briefly, one 0.5-ml straw of semen was diluted with 1.5 ml of a solution containing 10 mM TrisCl, 100 mM NaCl, 1 mM EDTA, pH 8.0) with 1% wt/vol sodium dodecyl sulfate, 1 mg proteinase K, and 40 mM dithiothreitol. This lysis solution was incubated overnight at 37°C in a 15 ml conical tube preloaded with 2 ml of high-vacuum grease in preparation organic phase extractions. The lysed and digested solution was extracted twice with 1 vol of phenol:chloroform:isoamyl alcohol (25:24:1), and once with 1 vol of chloroform. For each extraction the sample was centrifuged for 10 minutes in a swinging bucket rotor at 3210 x g at 23C to partition the organic phase below the band of high-vacuum grease, while to the aqueous phase was held above. The DNA was precipitated with 0.1 vol of 3 M sodium acetate (pH 5.2) and 2 vol of 100% ethanol. The precipitated DNA was washed once in 70% ethanol, briefly air dried, and dissolved in a solution of 10 mM TrisCl, 1 mM EDTA (pH 8.0). The bison tongue DNA was extracted with the Qiagen Blood and Cell Culture DNA Mini Kit according to the manufacturer’s instructions (Qiagen, Venlo, The Netherlands).

We sheared extracted DNA using a Covaris ultrasonicator (Covaris, Inc. Woburn, MA) prepared into Illumina sequencing libraries using either the NEB Ultra II FS kit (NEB, Ipswich, MA) or the TruSeq PCR-free DNA kit (Illumina, San Diego, CA). We quantified libraries with a Qubit fluorometer using the 1x HS kit (Thermo Fisher, Waltham, MA) or, for PCR-free libraries, via qPCR with the Kapa Complete Universal Kit (Roche Sequencing, Santa Clara, CA). Library fragment length distribution was assessed using either a Tapestation 2200 (Agilent, Santa Clara, CA) or a Fragment Analyzer (Advanced Analytical Technologies Inc., Ames, IA). For whole genome sequence (WGS), 1 μg of genomic DNA was fragmented and used to make indexed, 350 bp, paired-end libraries.

Beefalo and bison libraries were sequenced with Illumina instruments using 2 × 150 bp paired-end kits, either on the NovaSeq 6000 for bison or on the NextSeq 2000 platform for Beefalo and bison hybrids.

### Variant calling and genotyping

Remnant adapter sequences were trimmed from reads using Trimmomatic (v0.39) (Bolger, Lohse, and Usadel 2014), requiring a minimum length of 30bp. Trimmed reads were then mapped to the cattle genome ARS-UCD1.2 with the Y chromosome from Btau5.1 appended (ARS-UCD1.2_Btau5.0.1Y) using BWA (v0.7.17-r1188) mem (Li 2013) with default parameters, except for 9 Beefalo samples that were mapped with BWA aln (Li and Durbin 2009). These samples had a lower fraction of reads that were properly paired, with higher rates of interchromosomal mapping within read pairs, possibly suggesting chimera formation. As BWA aln conducts end-to-end alignment, it mitigates the effects of any spurious mapping of incompletely clipped chimeric sequences. Duplicate reads were removed with Picard MarkDuplicates (https://broadinstitute.github.io/picard/).

We called genotypes in four medium-to-high coverage (>10x) gaur genomes (Heaton et al. 2016; Verdugo et al. 2019; Wu et al. 2018) using GATK HaplotypeCaller (v4.1.8.1) (DePristo et al. 2011) to ascertain a set of variants from an outgroup to both cattle and bison for use in downstream analyses. Variants were filtered variants for a minimum genotype quality of 30 and minimum and maximum depths of 1/3rd and 2x the mean coverage on a per-sample basis. We also performed mappability filtering with SNPable (https://lh3lh3.users.sourceforge.net/snpable.shtml), using a *k*-mer length of 35 and stringency of 0.5 (gen_mask -l 35 -r 0.5).This yielded a set of 5,291,534 high-quality autosomal variants.

To accommodate the low sequencing depth we obtained from most Beefalo, we used psuedohaploid genotypes, in which a random read covering each gaur-ascertained variant was selected to represent genotypes at those sites. This approach mitigates biases arising from differential coverage across individuals (Barlow et al. 2020). Pseudohaploid genotypes were called for all individuals using PileupCaller using samtools (v1.9) mpileup (Li et al. 2009) with BAQ disabled (-B) and pileupCaller (https://github.com/stschiff/sequenceTools.git), requiring a minimum map quality of 25 and minimum base quality of 30.

### Modeling Beefalo ancestry

We visualized the relationships of Beefalo and bison hybrid individuals to bison and cattle using a Principal Component Analysis (PCA). PCA was conducted by computing principal components with *smartpca* (v18140) (Patterson, Price, and Reich 2006) using medium and high coverage bison, taurine cattle, and zebu cattle, and then projecting all Beefalo and bison hybrids onto these axes, using our set of pseudohaploid genotypes. This approach allows for directly understanding the ancestry of Beefalo individuals relative to these three groups while mitigating the effects of the low sequencing depth obtained for many Beefalo.

We used ADMIXTURE (v1.3.0) (Alexander, Novembre, and Lange 2009) to further model Beefalo ancestry. ADMIXTURE was run both in supervised and unsupervised modes, using the pseudohaploid dataset filtered for missingness (--geno 0.5) and minor allele frequency (--maf 0.05), and pruned for linkage disequilibrium (--indep-pairwise 50 5 0.5) using plink (v1.9) (Chang et al. 2015). Cross-validation was used to select the optimal value of K. For the supervised ADMIXTURE analysis, Beefalo and bison hybrid ancestry was fit as potentially coming from bison, taurine, or zebu sources.

We also sought to investigate Beefalo ancestry on the sex chromosomes. For the X chromosome, there were 649,441 total variants, which were filtered to require that they were called in 66 individuals and had a minor allele frequency of at least 0.05, with 97,198 variants remaining. We used the 93 males within the overall set of samples with a level of missingness less than 25% and ran ADMIXTURE in haploid mode. We estimated the Y chromosomal phylogeny using males in the sample set with several additional species added as outgroups, including yaks, European bison, gaur, and banteng. The phylogeny was estimated from 18,056 variants with IQ-TREE (1.6.12) (Nguyen et al. 2015), using the GTR+ASC model and obtaining branch support values using ultrafast bootstrapping (Hoang et al. 2018).

After exploring overall autosomal Beefalo ancestry using model-free (PCA) and model-based (ADMIXTURE) approaches, we sought to specifically test for the presence of bison ancestry in Beefalo and bison hybrid genomes. The D-statistic *D(Beefalo, taurus; bison, water buffalo)* was used for this, which tests for excess allele sharing between Beefalo and bison, relative to taurine cattle, with water buffalo as an outgroup. We had observed that Beefalo formed a cline between taurine and zebu cattle in PCA, so we also used the statistic *D(Beefalo, taurus; indicus, water buffalo)* to test for zebu ancestry among individual Beefalo. Finally, we quantified the proportion of bison ancestry in Beefalo genomes using the f4-ratio *f4(yak, water buffalo; Beefalo, taurine)/f4(yak, water buffalo; bison, taurine)*, which estimates the proportion of ancestry, α, in Beefalo that comes from lineage related to bison, relative to a yak outgroup (the closest outgroup to bison). D- and *f-*statistics were calculated using ADMIXTOOLS2 (Maier et al. 2023) for each individual Beefalo and bison hybrid, grouping bison and taurine and zebu cattle.

### Locating segments of differential ancestry within Beefalo

Diploid ancestry state across individual Beefalo genomes was modeled as coming from bison and cattle-related sources by using AncestryHMM (v1.0.2) (Corbett-Detig and Nielsen 2017). We used medium-to-high coverage genomes (>10x) from bison (n=20), taurine cattle (n=20), and zebu (n=6) each as source panels. We called genotypes for each species separately using the same filtering procedure as with our outgroup-ascertained variants. We then merged these panels, requiring sites to be called in at least 6 bison, 6 taurine cattle, and 3 zebu, have a frequency difference of at least 90% between any two source panels, and be at least 500bp from the nearest variant. This gave a final set of 2,322,535 ancestry informative markers. We modeled local ancestry across the genome for each Beefalo using AncestryHMM with the parameters ’-e 0.02 -a 3 0.02 0.90 0.08 -p 0 -5 0.02 -p 1 10000 0.98 -p 2 -20 0.08’. This models two gene flow events, one 20 generations ago from zebu cattle at 8% and the second from bison 5 generations ago at 2%, though allows the exact timing of these admixture pulses to be inferred. We used read counts at ancestry informative markers for all Beefalo as input for ancestry inference. For seven individuals which were sequenced to high coverage, we performed additional ancestry inference using both read data downsampled to ∼2x coverage and genotypes at ancestry informative positions. Genotypes at ancestry informative markers were called using GATK HaplotypeCaller. We filtered posterior ancestry probabilities using a 0.9 threshold.

## Tables

Supplementary Table 1: List of samples used

## Acknowledgements

We thank USMARC technicians H.R. Sadd and N.J. Allison for assistance with laboratory procedures. Mention of trade names or commercial products in this publication is solely for the purpose of providing specific information and does not imply recommendation or endorsement by the USDA. The USDA is an equal opportunity provider and employer.

## Data Availability

Beefalo, bison hybrid, and bison sequencing data generated in this study are available in NCBI BioProject PRJNA1152308.

## Supplementary Figures

**Supplementary Figure S1.**
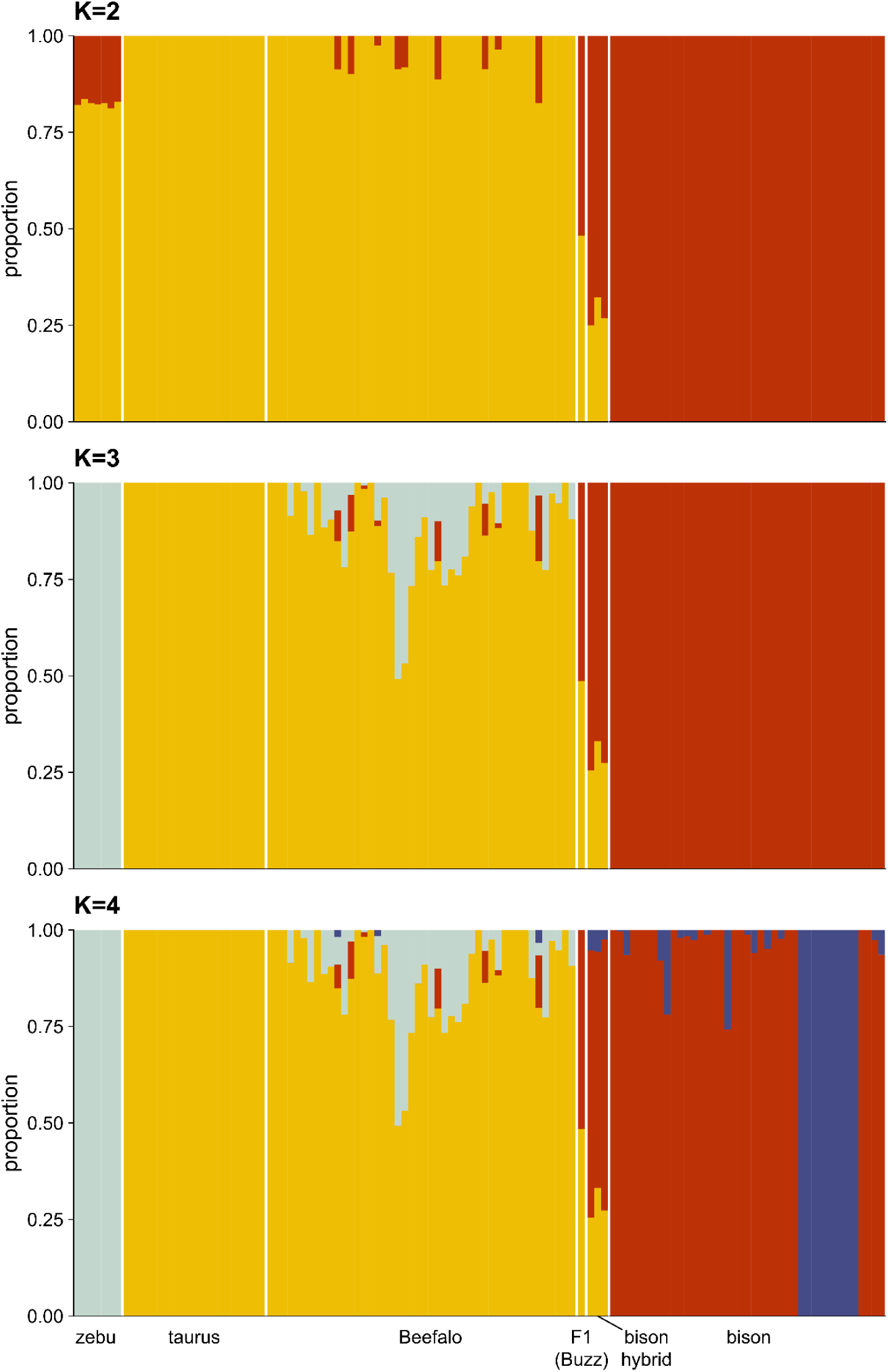
Unsupervised ADMIXTURE analysis of cattle, bison, bison-cattle hybrids, and Beefalo at different values of K, from K=2 to K=4. At K=4, bison are split into two groups, which correspond to bison subspecies (wood and plains bison).

**Supplementary Figure S2.**
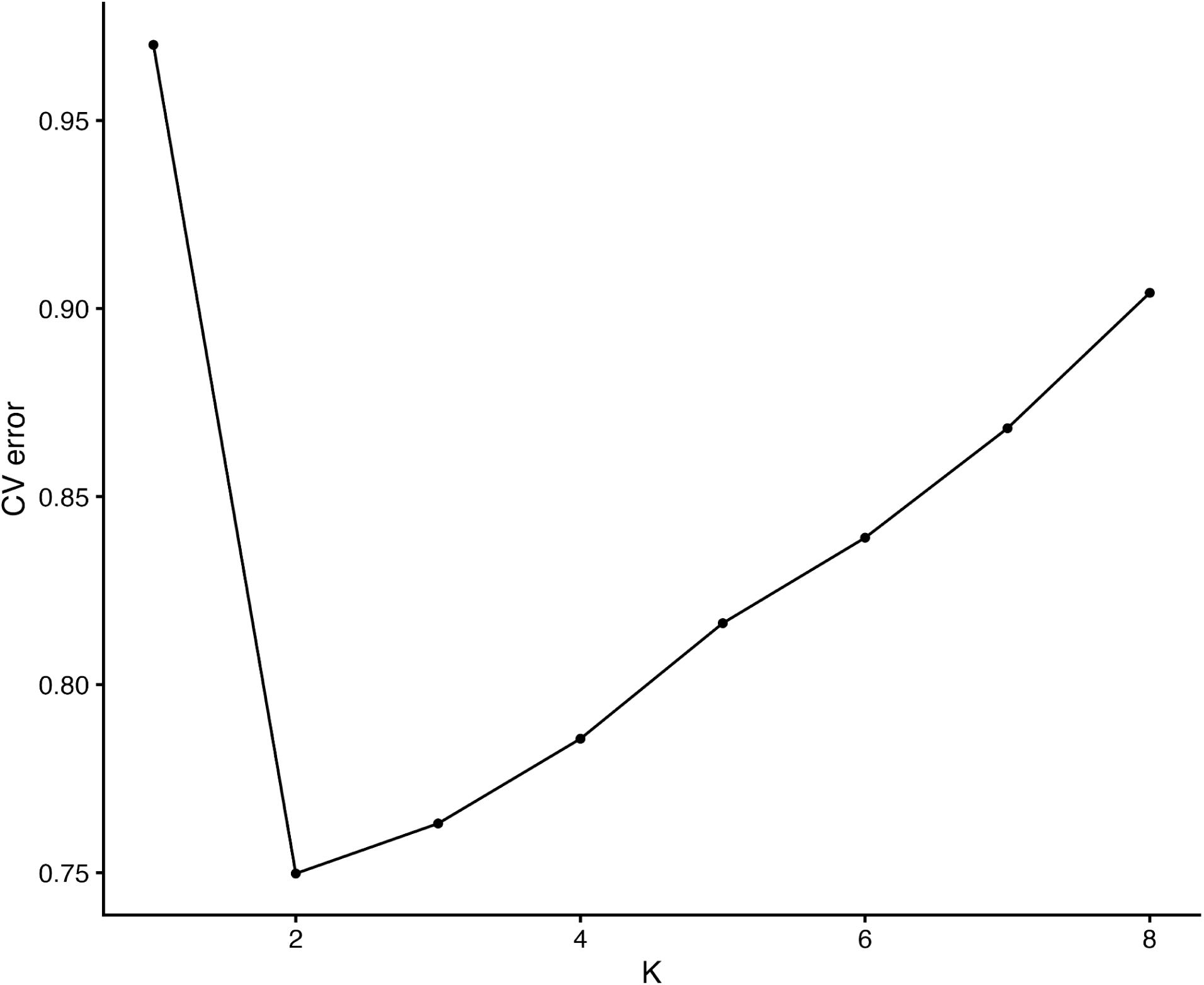
Cross-validation results comparing unsupervised ADMIXTURE using cattle, bison, bison-cattle hybrids, and Beefalo across different values of K, ranging from 1 to 8.

**Supplementary Figure S3.**
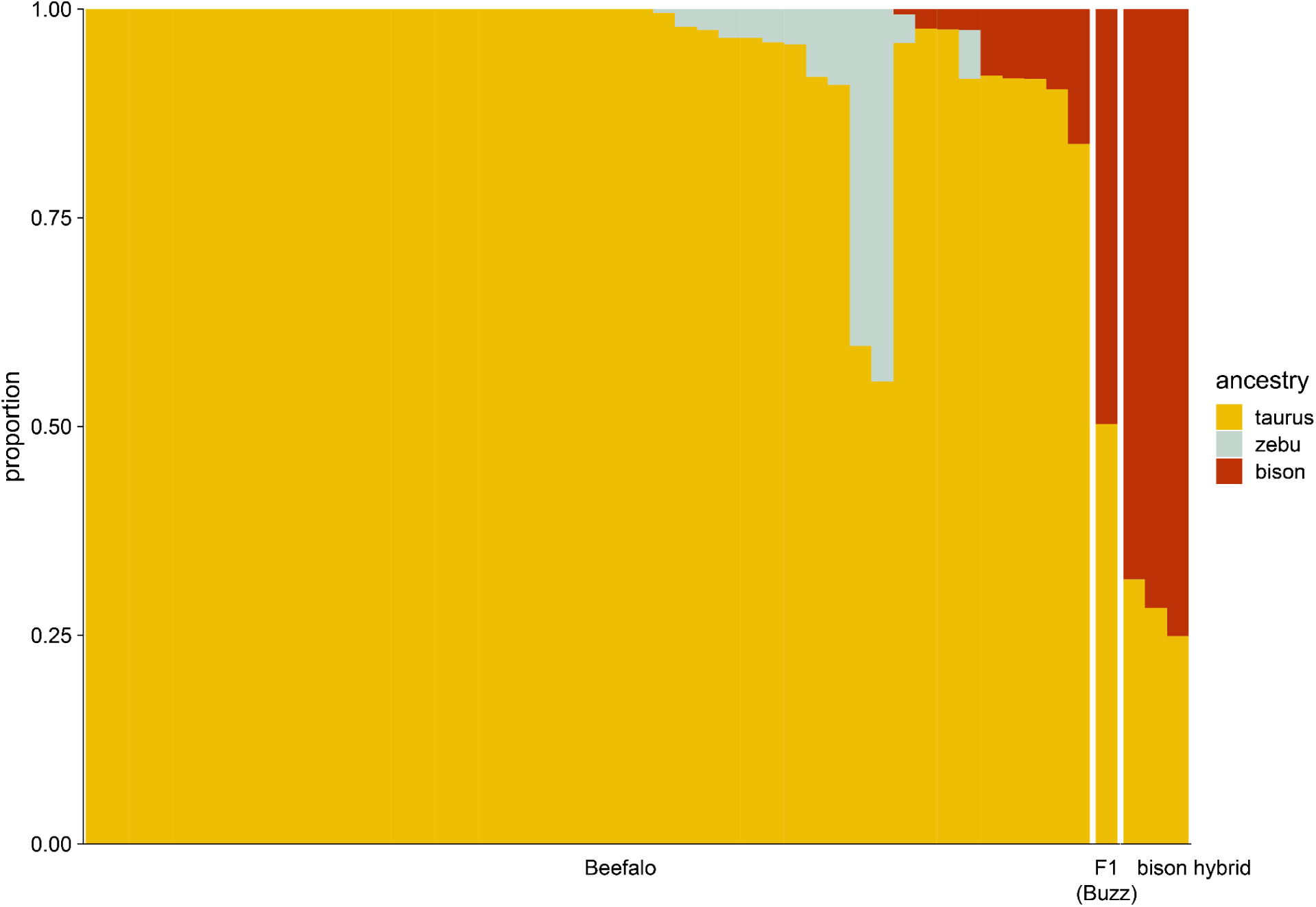
Supervised ADMIXTURE modeling of Beefalo and bison hybrid ancestry, using panels of bison and taurine and zebu cattle.

**Supplementary Figure S4.**
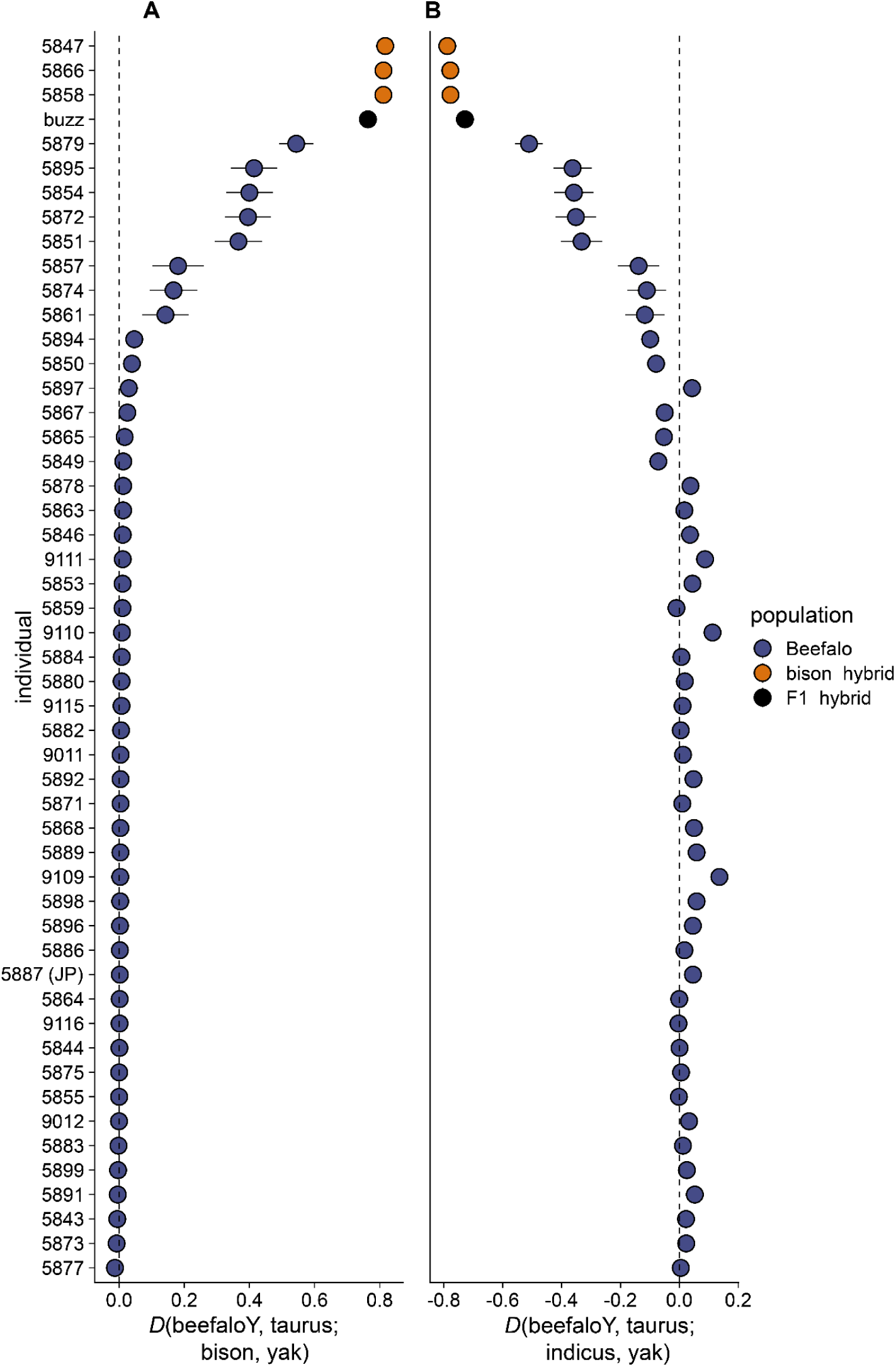
Allele sharing statistics using individual bison hybrids and Beefalo. This is the same as Fig. 2, except yak is used as an outgroup instead of water buffalo. **A)** D-statistics testing for allele sharing between individual Beefalo and bison hybrids, relative to taurine cattle. **B)** D-statistics testing for allele sharing between Beefalo and zebu cattle, relative to taurine cattle. For all panels, error bars depict 3 standard deviations.

**Supplementary Figure S5.**
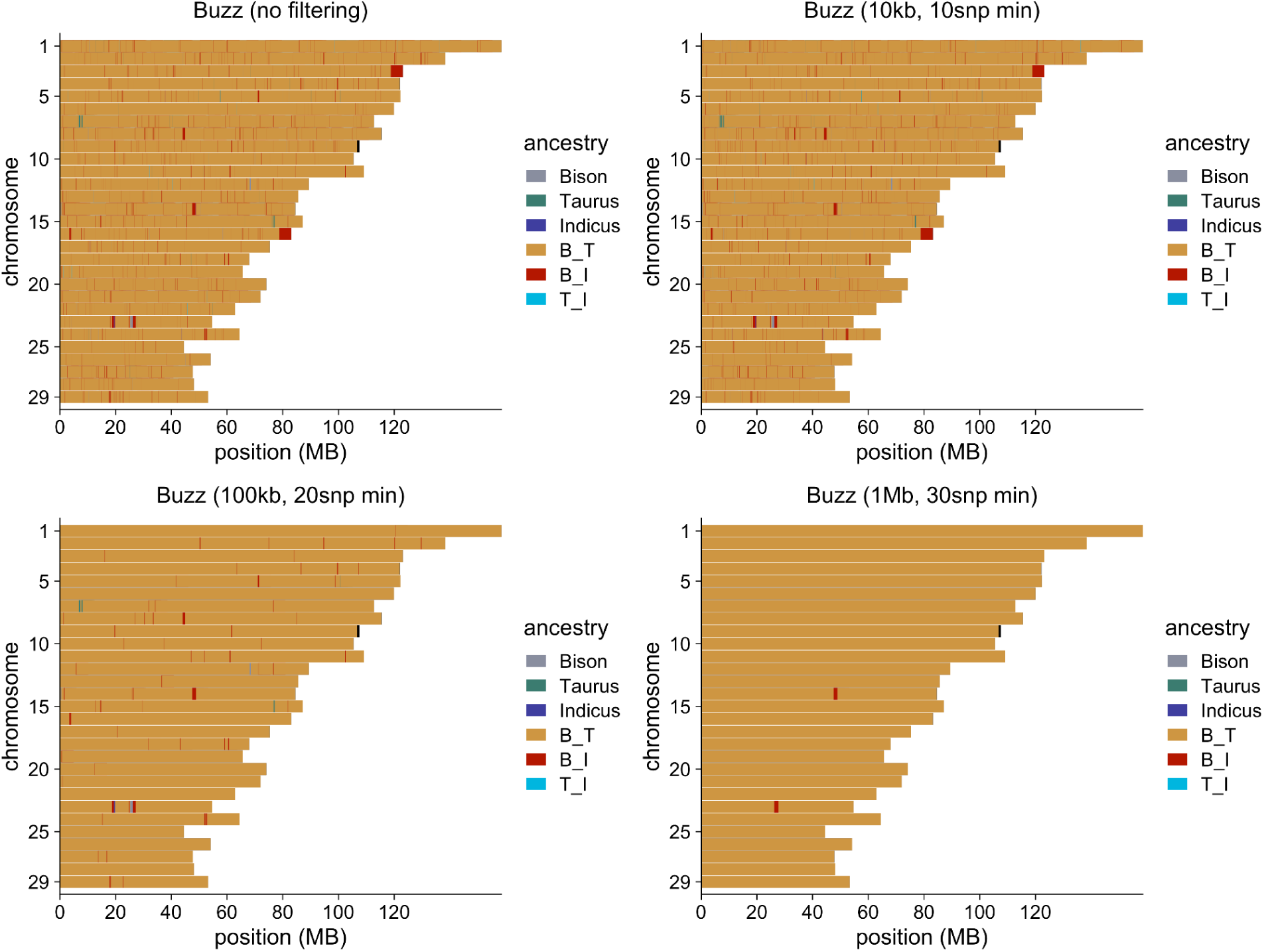
Local ancestry inference results from a F1 bison-taurine cattle hybrid. Ancestry was correctly inferred to be almost completely heterozygous for bison and taurine ancestry across the genome, though with some small segments incorrectly assigned to be heterozygous for bison-zebu bison, likely because of the low divergence between cattle subspecies. These segments are all small and are increasingly removed with more stringent filtering (for window size and minimum SNPs to call a window, shown in panel titles).

**Supplementary Figure S6.**
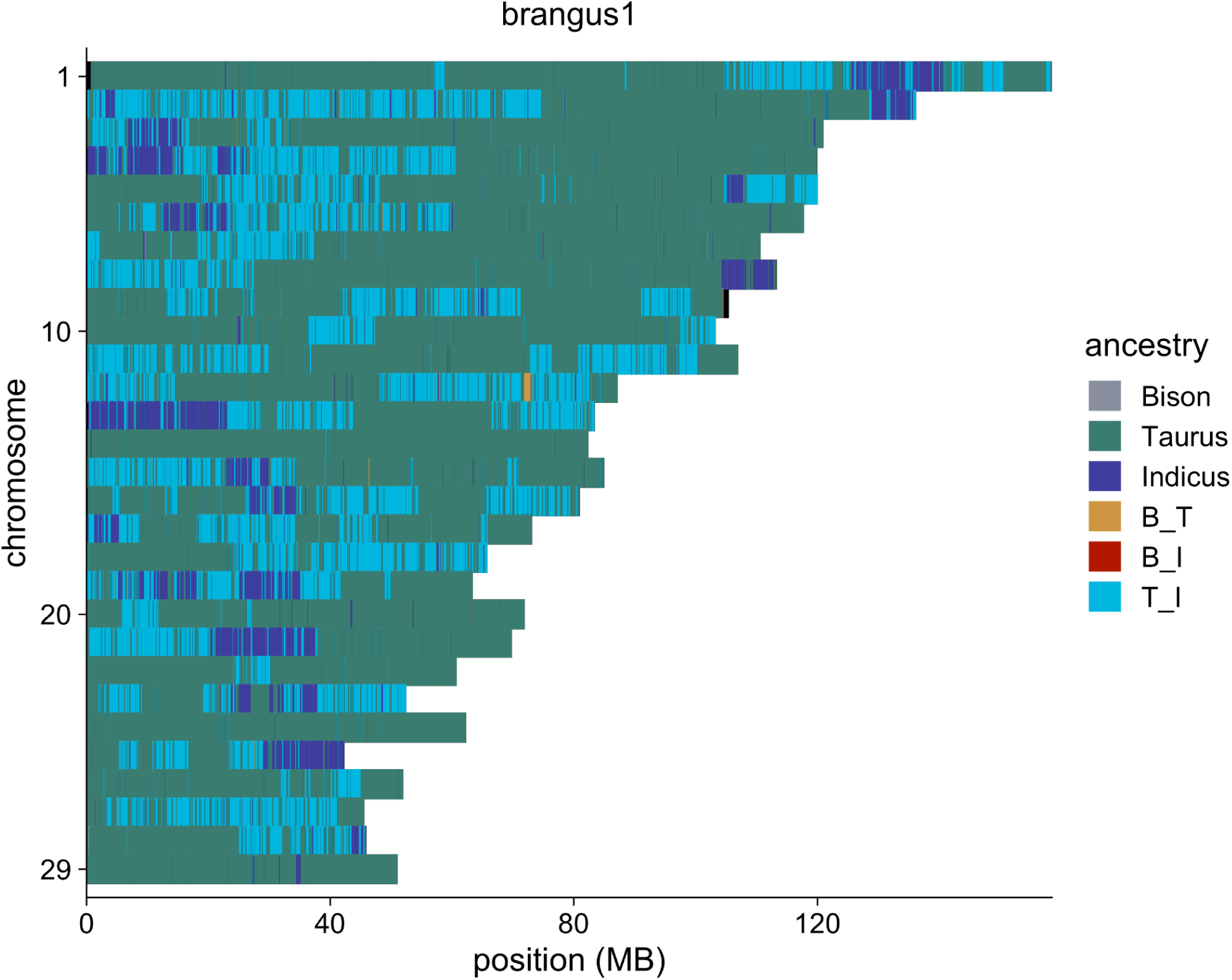
Local ancestry inference results from a Brangus individual, showing the presence of both taurine and zebu (indicine) ancestry.

**Supplementary Figure S7.**
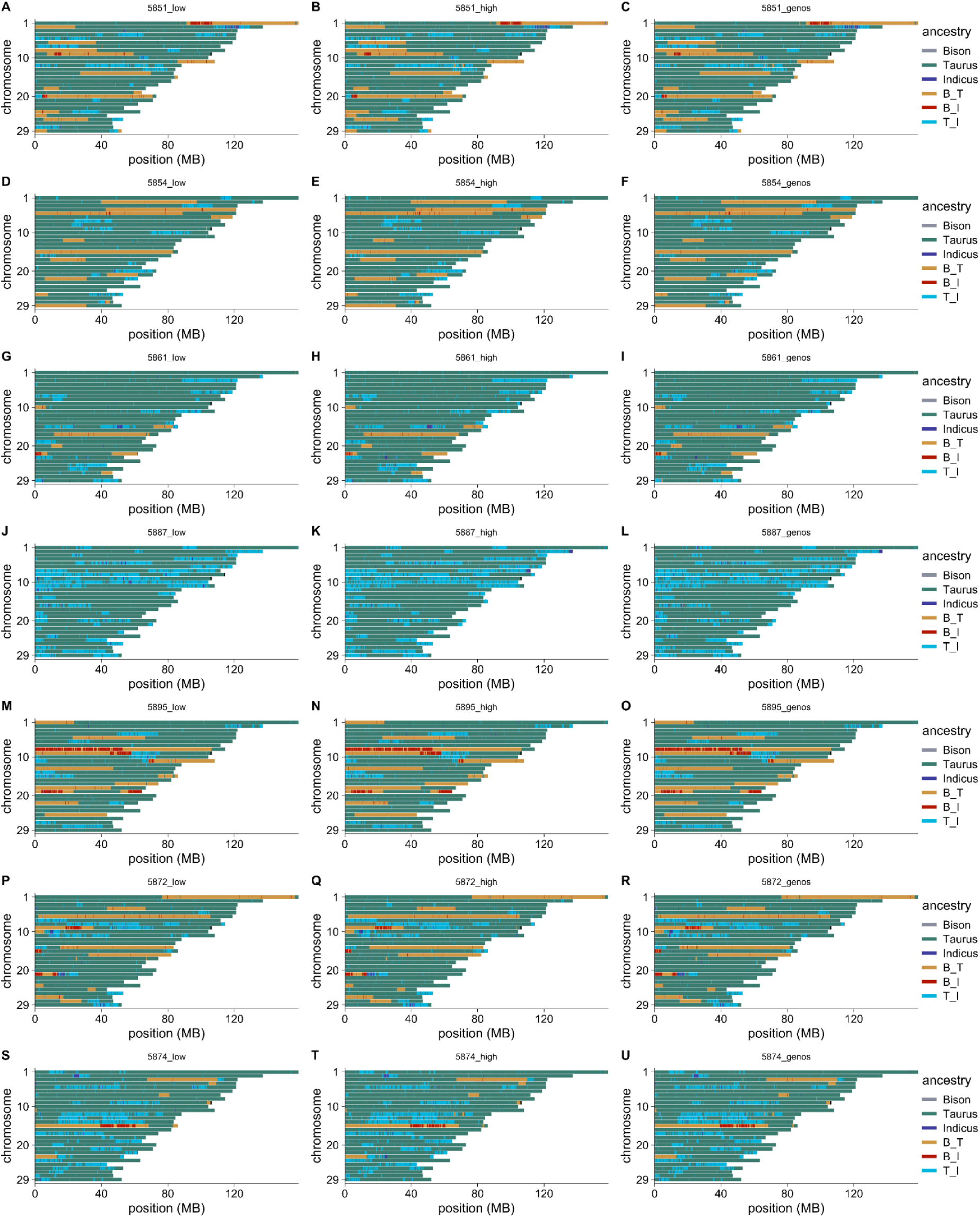
Comparison of local ancestry inference using either downsampled (∼2x) read data (A, D, J, M, P, S), high coverage (30-42x) read data (B, E, H, K, N, Q, T), or called genotypes (C, F, L, O, R, U) on seven individual Beefalo and bison hybrids which were sequenced to high coverage: 5851 (A-C), 5854 (D-F), 5861 (G-I), 5887 (J-L), 5895 (M-O), 5872 (P-R), 5874 (S-U).

## Notes

### Competing Interest Statement

The authors have declared no competing interest.

### Summary of Updates

This manuscript has primarily been revised to provide additional quantitative detail throughout the main text, including the range of Z scores obtained from allele sharing tests, the exact number of Beefalo individuals detected with either bison or zebu cattle ancestry, and the range of estimated ancestry proportions. PCAs now include the proportion of variance explained for each axis, and we more consistently show the reported bison ancestry content of Beefalo individuals throughout the figures. ADMIXTURE results for K=2 and K=4 have been added as a supplemental figure, as well as a figure depicting ADMIXTURE cross validation results from K=1-8 to support the choice of K=3 for the main text. Additionally, we clarified how outgroups were defined and used in our study and have included a supplementary figure showing allele sharing results using yak as an outgroup, instead of water buffalo as in the main text. We also added further quantitative detail to the methods, such as the number of SNPs used in our analyses, and emphasized our outgroup SNP ascertainment and pseudohaploid genotype calling approaches for generating the dataset used for analyses. Additionally, some aspects of the history of the Beefalo breed have been clarified and we have added additional details about how our sampling scheme is sufficient for surveying bison ancestry across the breed. Some additional information about zebu cattle breeds has also been included, and this group of cattle is now referred to exclusively as zebu, instead of with a mix of zebu and indicine.

### Funding

- Howard Hughes Medical Institute: Investigator Award to BS

